# A high-speed, modular display system for diverse neuroscience applications

**DOI:** 10.1101/2022.08.02.502550

**Authors:** Matthew Isaacson, Lisa Ferguson, Frank Loesche, Ishani Ganguly, Jim Chen, Andy Chiu, Jinyang Liu, William Dickson, Michael Reiser

**Affiliations:** Howard Hughes Medical Institute, Janelia Research Campus, Ashburn, VA; Meinig School of Biomedical Engineering, Cornell University, Ithaca, NY; Center for Theoretical Neuroscience, Columbia University, New York, NY; Sciotex, Newtown Square, PA; California Institute of Technology, Pasadena, CA & IO Rodeo Inc. Pasadena, CA

## Abstract

Visual stimulation of animals in the laboratory is a powerful technique for studying sensory control of complex behaviors. Since commercial displays are optimized for human vision, we established a novel display system based on custom-built modular LED panels that provides millisecond refresh, precise synchronization, customizable color combinations, and varied display configurations. This system simplifies challenging experiments. With variants of this display, we probed the speed limits of motion vision and examined the role of color vision in behavioral experiments with tethered flying *Drosophila*. Using 2-photon calcium imaging, we comprehensively mapped the tuning of visual projection neurons across the fly’s field of view. Finally, using real-time behavior analysis, we developed low-latency interactive virtual environments and found that flying flies can independently control their navigation along two dimensions. This display system uniquely addresses most technical challenges of small animal vision experiments and is thoroughly documented for replicability.

## Introduction

Visual systems are a major entry point to studying brain function across animals. The study of insect vision has been critical for understanding neural mechanisms of sensory processing and the role of vision in organizing behavior^1,2^. Mechanistic studies of vision require precise control of an animal’s visual experience, which is often accomplished by immobilizing the animal’s head and/or body while presenting visual stimuli. Early studies in insects used rotating cylinders lined with different patterns^3,4^. Variations of this approach relying on physical movement enabled studies of behavioral and neuronal responses to movement along a single-degree of freedom, usually rotation^5^, but are not well suited to displaying more complex visual stimuli. An early innovation used patterned light projected onto screens in front of restrained animals^1,6,7^. In recent years, advances in computing and display technologies have enabled a range of alternative experimental options, including displays based on light-emitting diodes (LEDs)^8–11^. Even more specialized display systems have been created to fill specific experimental needs. For example, the pioneering work characterizing the motion sensitive neurons in flies relied on an ingenious, rotatable mechanical stage that could present local motion of a rotating visual object over a large visual area^12^. Since no display method is ideal for all experiments, we developed displays constructed from LED ‘panel’ modules^13^ that enabled numerous experiments, such as visual place learning^14^, landmark orientation^15^, fine-scale mapping of neuronal receptive fields^16^, and multi-sensory integration^17^. Recently the system has been adapted to build a spherical arena for stimulating the entire visual field of larval zebrafish^18^. While this system has clearly been useful, many experiments would benefit from improvements in the complexity, resolution, and speed of visual stimuli, and from more flexible integration into diverse experimental systems.

Like nearly all electronic technology, commercial LED-based display systems have improved considerably in recent years, but these systems are designed for human use, and therefore are not ideal for many animal experiments. Displays are typically built from red, green, and blue LEDs to match the spectral sensitivity of human cone cells^19^, but most animals process the visual world using photoreceptors with different spectral sensitivity^20^. Importantly, the most popular genetic model organisms use in neuroscience research—fruit flies, mice, and zebrafish—all see with UV-sensitive photoreceptors^21–23^, suggesting that commercial display systems will never be suited for studying color vision in these animals. Most displays developed for humans refresh at 60 Hz, which is below the flicker fusion frequency of many insect eyes^24^, and leaves little headroom for accurately displaying fast, apparent motion stimuli. Commercial displays prioritize smooth presentation for viewers relying on frame buffers, which often means there is a significant latency between received and displayed information, complicating experiments that require precise timing of stimuli and synchronization with other experimental events. Consumer displays are often large, flat rectangles, an adequate design for humans with forward-facing eyes and prominent, central foveae. However, these displays only cover small regions of the visual field of animals with side-facing eyes and limited foveal vision. More panoramic stimulation can be achieved by arranging multiple rectangular displays^25^, by projecting onto multiple flat screens^26^, or a single large toroidal screen^27^. These approaches have some drawbacks for vision research beyond increased setup complexity, especially non-uniform brightness and contrast across the display. While it is an exciting time for visual neuroscience with many available display solutions, the myriad tradeoffs and limitations suggest that a more advanced, yet general solution could improve the quality and precision of experiments in animal vision.

The modular LED display system described previously^13^ (and referred to as “G3” for 3^rd^ generation) has received many incremental updates to improve its functionality, although it is based on a ∼15-year-old design. We have developed and validated the “G4” (4^th^ generation) modular display system that improves on all aspects of the prior design. G4 includes modern technological advances and a simplified, yet more integrated experimental control system. In this paper, we describe the hardware architecture and design strategy of the modular G4 system (Figure 1) that enables full-display brightness control at 1 kHz refresh rates, and precise synchronization with external equipment. We then introduce new software tools that we developed to simplify the use of a G4 display for end users (Figure 2). We summarize our thorough validation of the system, first using technical benchmarks, and then using different G4 configurations in novel experiments that probe behavioral and neural responses of *Drosophila*. Each of the experiments takes advantage of unique, advanced features of the G4 system. The fast display refresh is used to characterize the upper limits of motion detection in flies across a variety of self-motion types (Figure 3). Flexible integration with external equipment enables a motor-controlled display that we used to map the local motion sensitivity of visual projection neurons across the fly’s field of view (Figure 4). Fine brightness control of a UV/green G4 display was used to demonstrate that moving patterns in each spectral channel can be perceptually balanced, demonstrating isoluminance (Figure 5). Finally, using fast, closed-loop simulation of object tracking, we measured the effects of display latency on *Drosophila* flight behaviors, and we demonstrated closed-loop behavior control strategies in complex virtual environments (Figure 6).

**Figure 1:**
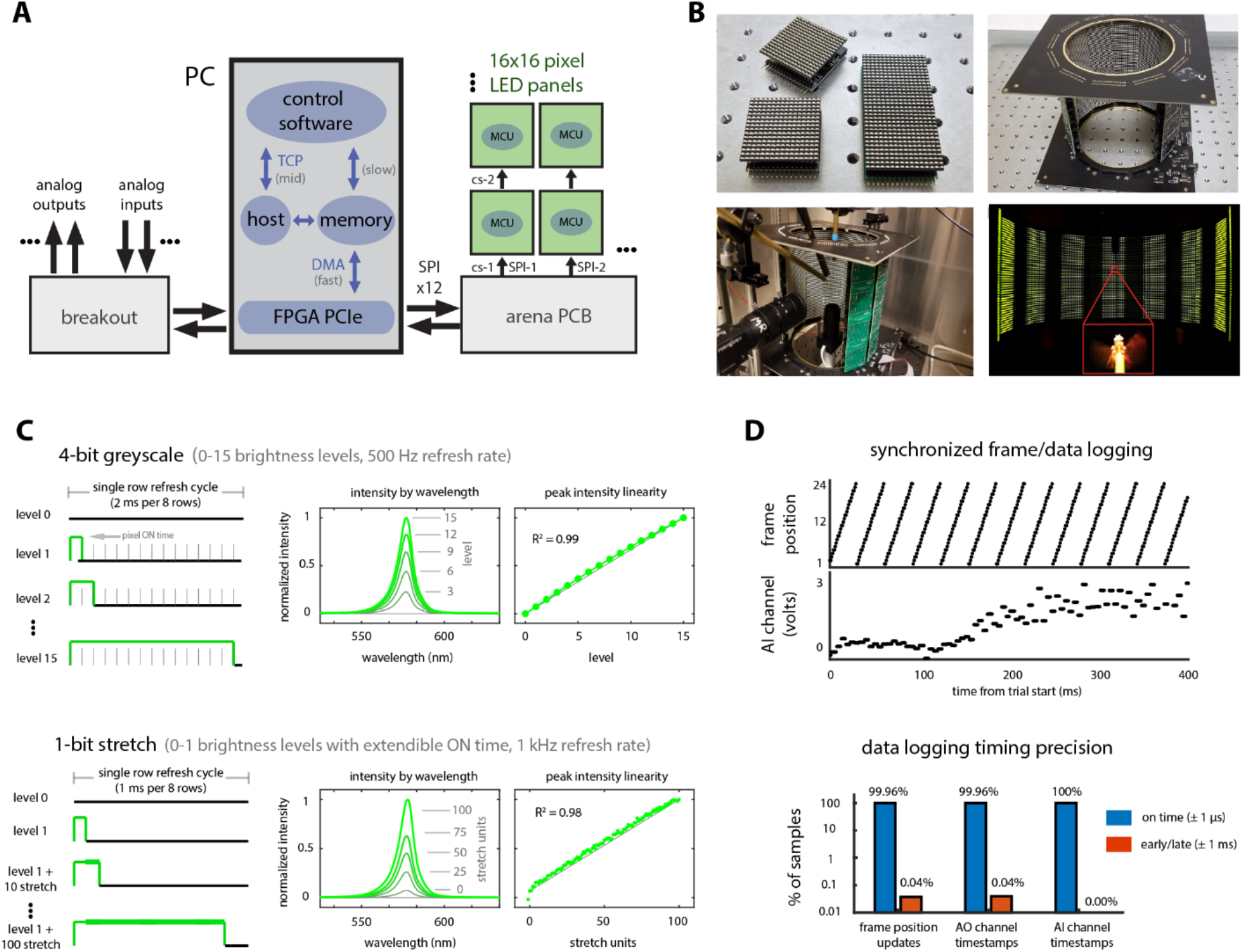
The G4 modular display system. A) Modular LED display system architecture. On the PC, high-level control software generates and stores patterns to PC memory and sends commands to the host application over TCP/IP. The host directs the FPGA to access stored pattern data via a fast Direct Memory Access (DMA) connection (<1 ms latency). The FPGA streams pattern data over 12 parallel SPI busses to the LED panels of a display arena, simultaneously logging analog inputs and controlling analog outputs connected through an external breakout box. B) Images of an experimental setup with a G4 display, assembled from individual 40 mm × 40 mm panels. Cylindrical arenas are assembled with multiple columns of LED panels, delivering a panoramic visual stimulus to a tethered, flying fly. C) Displaying 1-bit or 4-bit patterns with 2 or 16 brightness levels respectively. For 4-bit patterns (top), new frames are refreshed at 500 Hz through row-scanning (8 rows per quadrant refresh cycle), and pixel-by-pixel brightness is controlled using LED ON pulses of 0-15 duration levels, achieving linear brightness control. (left) Schematic of the ON pulses for example brightness levels 0, 1, 2, and 15. Spectra (middle) and total luminance (right), normalized to the maximum brightness (level 15), of the green LEDs for the indicated brightness levels. For 1-bit patterns (bottom), the LEDs are refreshed at 1 kHz, and global brightness modulation is controlled by increasing the duration of the ON pulse (in “stretch” units) of all 8 rows, on a frame-by-frame basis. Normalized spectra and overall brightness are shown for the indicated brightness levels/stretch units. D) Precise synchronization of analog inputs/outputs to the display update cycles. (top) Example experiment data showing ms-precision synchronization between the display refresh cycles of a 24-frame drifting grating pattern (rightward yaw rotation) looping at a temporal frequency of 32 Hz, synchronized to the analog input channel (ADC0) acquiring a flying fly’s turning reaction to the grating stimulus (0 = no turn, positive values = right turn). (bottom) Distribution of timing errors between desired vs. actual timestamps of logged analog inputs (ADCs) and analog outputs (AOs) synchronized to display refresh cycles, with errors categorized as “on-time” (i.e. within 1 µs of the intended time) or “early/late” (between 1 µs and 1 ms from the intended time).

**Figure 2:**
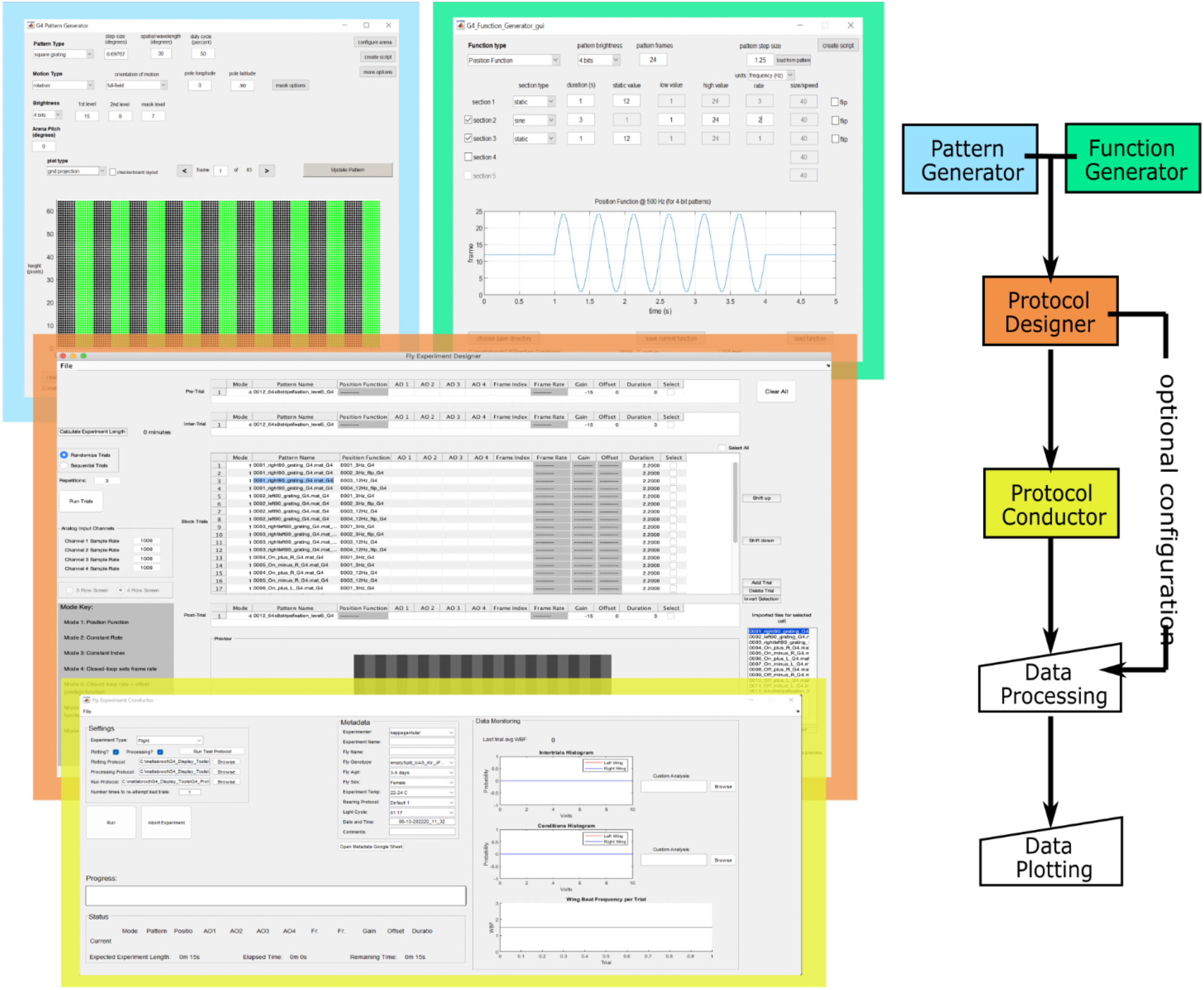
Software tools for the G4 display system. Flowchart of MATLAB software tools (right) and screenshots of the Graphical Users Interfaces (GUIs, on left) for designing, running, and analyzing experiments. Patterns (e.g. drifting gratings, edges, starfields) are generated from parameterized settings with the Pattern Generator GUI and/or scripts (blue). Position functions, which control the timing for displaying a pattern’s frames on the display, and analog output functions, which control voltage signals on the system’s 4 analog outputs, are similarly created using the Function Generator (green). The Protocol Designer (orange) GUI is used for creating experiments that control the timing and order in which pre-generated patterns and functions are displayed, as well as how the data generated from the experiment will be processed afterwards; protocols can be saved and run later or executed immediately. Protocols are executed using the Protocol Conductor GUI (yellow), which logs experiment data (and metadata from the user) while updating the current state and overall progress of the experiment in real-time. Streamed analog input data can be plotted in the Conductor GUI after each trial for additional monitoring. Raw experiment data and metadata files are then processed into easily accessible data structures using the Data Processing scripts and visualized using Data Plotting scripts (white). Depending on the configuration in the Protocol Designer, these scripts can be triggered by the Experiment Conductor or run manually. These tools enable end-users to design and conduct highly reliable experiments with precisely specified timing, without writing any code.

**Figure 3:**
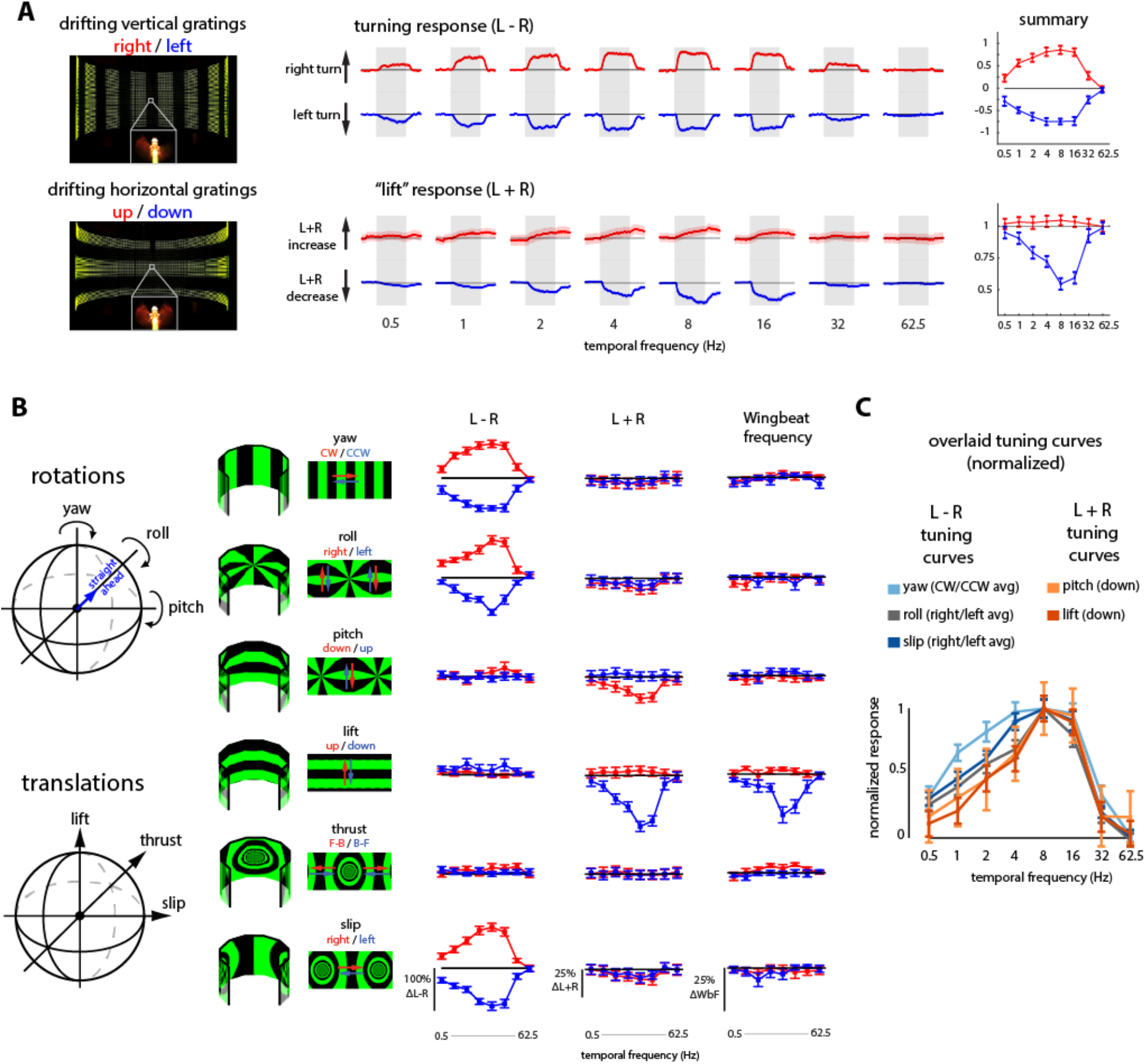
Determining the upper limits of *Drosophila* motion vision using high speed optic flow stimuli. A) Example snapshots (left) of a tethered flying fly’s reactions to horizontally drifting vertical gratings (“yaw” motion, top) and to vertically drifting horizontal gratings (“lift” motion, bottom). Complete sequences shown in Movie S1. Time-series of the turning response (middle, top), measured as the difference in wingbeat amplitudes (L-R) in response to clockwise and counter-clockwise yaw motion, and the lift response (middle, bottom) measured as the sum of wingbeat amplitudes (L+R), to gratings drifting up and down. The gray shaded region demarcates the 2 s interval during which the stimulus is presented at the indicated speeds. The time-series are plotted as mean ± SEM from N=21 flies. These data are summarized as temporal frequency tuning curves on the right (mean ± SEM). B) Flight behavioral reactions to grating patterns designed to present 6 types of optic flow: 3 rotations and 3 translations, each drifted in both directions. The patterns are displayed in both a 3D cylindrical view and an “unwrapped” view (left 3 columns). Temporal frequency tuning curve summaries of L-R, L+R, and wingbeat frequency are shown for all motion patterns. Data are presented as mean ± SEM from the same N=21 flies and include the data in (A) (right 3 columns). C) The normalized tuning curves of turning responses across motion patterns (except for thrust motion, shown in Figure S1). Symmetrical L-R reactions to both directions of yaw, roll, and slip rotations are averaged together (after sign correction), while for asymmetric L+R reactions to pitch and lift translations, only the larger reactions to downward motion are shown.

**Figure 4:**
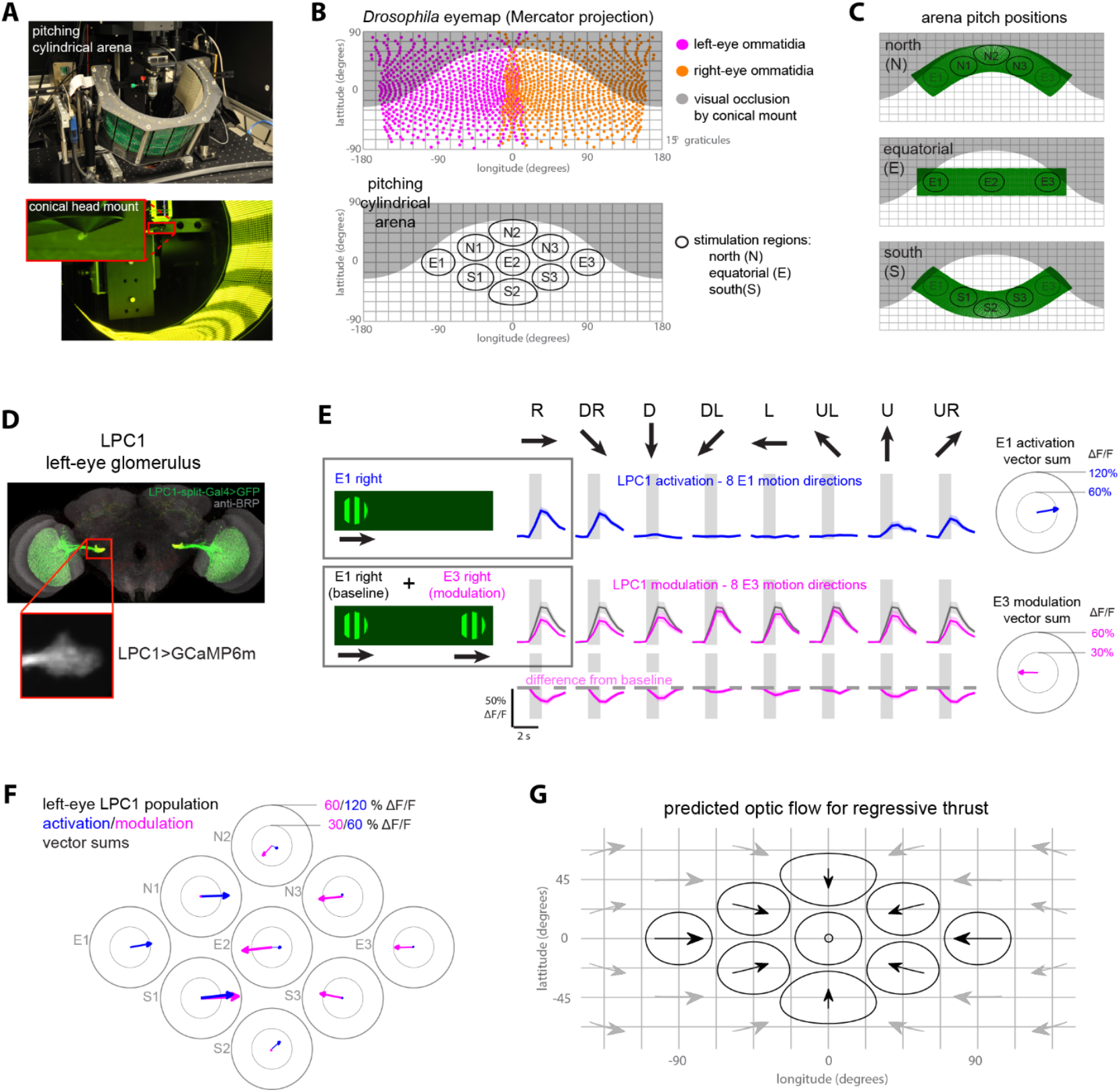
Mapping the directional tuning of visual projection neurons using a motorized visual display. A) Pictures of 2-photon imaging setup using a low-occlusion conical head mount and motorized, pitching cylindrical arena. B) Mercator projection estimate of the *Drosophila* visual field (top) using optical axes previously reported^56^. Visible area in white, occluded area in gray (new design compared to prior design in Figure S2). Spatial locations targeted for receptive field mapping of the fly’s un-occluded visual field (bottom). C) The spatial locations (in Mercator projection) of the pitching LED arena used to stimulate the 9 receptive field locations: north for presenting motion in spots N1-3; equatorial for E1-3; south for S1-3. D) The split-GAL4 line used to target LPC1 neurons. Confocal stack of the driver line expressing GFP (top), below is a typical 2-photon imaging field of view of the left LPC1 glomerulus expressing GcaMP6m. E) LPC1 activation and modulation in response to motion stimuli in 8 directions in spots E1 and E3. Mean LPC1 activation (top row) by motion along the indicated direction (square-wave gratings, 100% contrast, 8 Hz temporal frequency) presented in location E1 (ipsilateral to imaged LPC1 glomerulus). Example stimulus for E1 right is shown. Mean modulation of the LPC1 response to E1 rightward motion induced by simultaneously presenting motion stimuli in location E3 (contralateral to the LPC1 glomerulus). Baseline response to E1 rightward motion in gray; modulated response by simultaneous E3 motion in magenta. Example stimulus for E3 right is shown. Data plotted as mean ± SEM, N=8 flies. The combined vector sum (right) of responses (E1 activation on top, E3 modulation on bottom) from all flies to 8 motion directions. Individual response vectors were calculated as the product of each motion direction and response amplitude. F) (left) Combined vector sums of LPC1 activation (8 directions at 9 spatial locations) and modulation (8 directions at 8 spatial locations; E1 is reserved for baseline activation). Individual fly data from the N=8 flies in Figure S3. (G) Predicted directions of apparent motion across the visual field due to regressive thrust translation.

**Figure 5:**
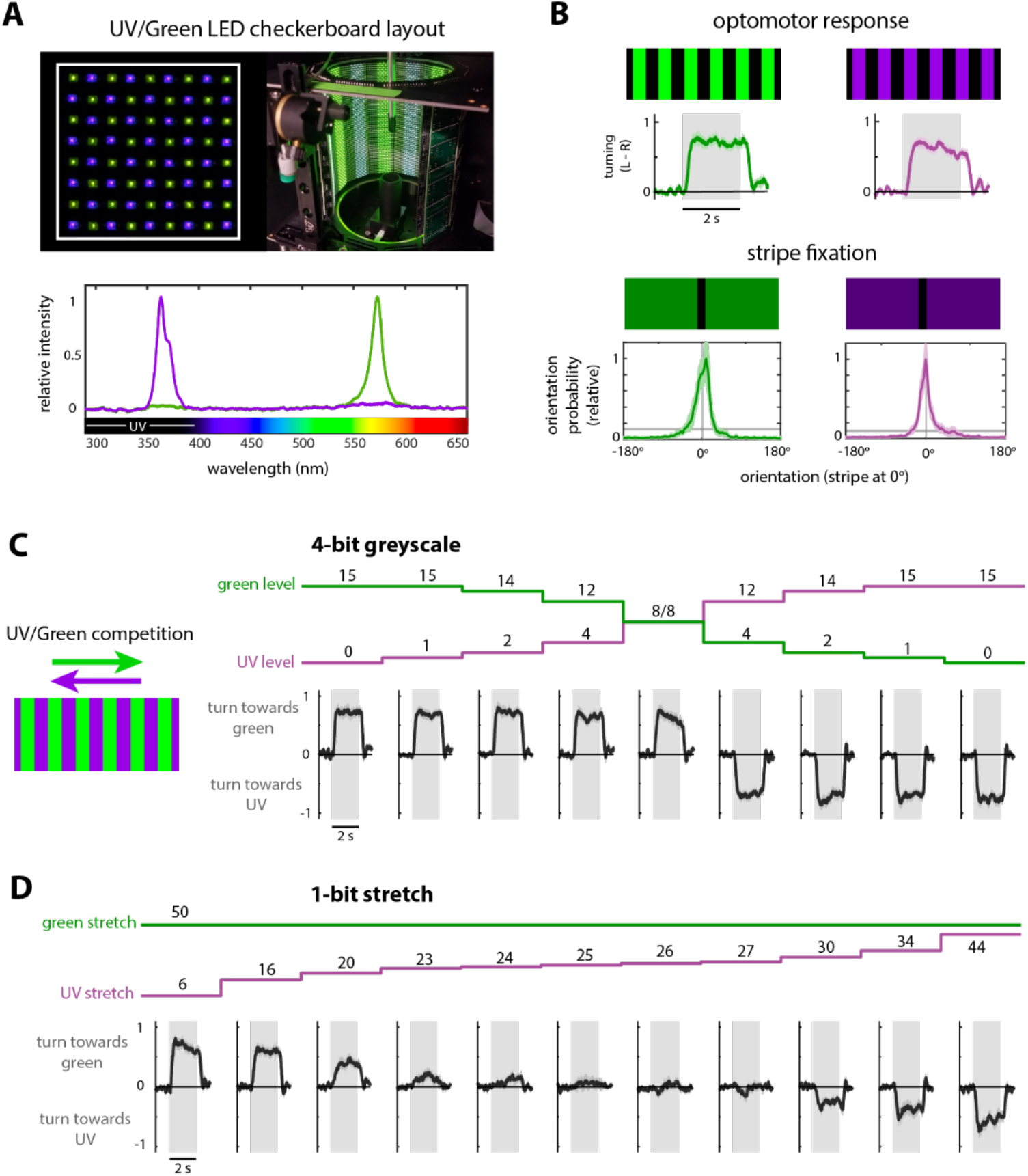
Exploring the interactions between UV and green visual motion responses using precise brightness control. A) 16×16 LED panel composed of UV and green LEDs in a checkerboard layout (top, left) was used to assemble a cylindrical arena (top, right) for 2-color visual stimulation. Spectra (bottom) of the UV and green LEDs (normalized to peak of each) in the 2-color panels. B) Time-series turning (L-R) responses of tethered, flying flies (top) to clockwise rotation of a grating pattern (alternating bars of brightness levels 0 and 15, 8 Hz temporal frequency) displayed with either only green (left) or only UV (right) LEDs in the 2-color arena. Histogram of flies’ orientation (bottom) during closed-loop stripe fixation with a dark vertical bar (brightness level 0, 30° width) on a medium brightness (level 8, UV or green) background. C) Normalized average turning responses (L-R) of flying flies to competing UV/green grating stimuli composed of overlaid green and UV gratings simultaneously drifting in opposite directions. The patterns with 16 brightness levels were used to independently control the brightness of the UV and green bars. D) Turning responses to competing UV/green gratings, where 2 level (on/off), “stretch” modulation was used to control the relative UV/green brightness levels by stretching the on LED time, to establish conditions of isoluminance, where the motion of the patterns is apparently balanced for the flies, resulting in no net turning. All data are from N = 8 flies, plotted as mean ± SEM.

**Figure 6:**
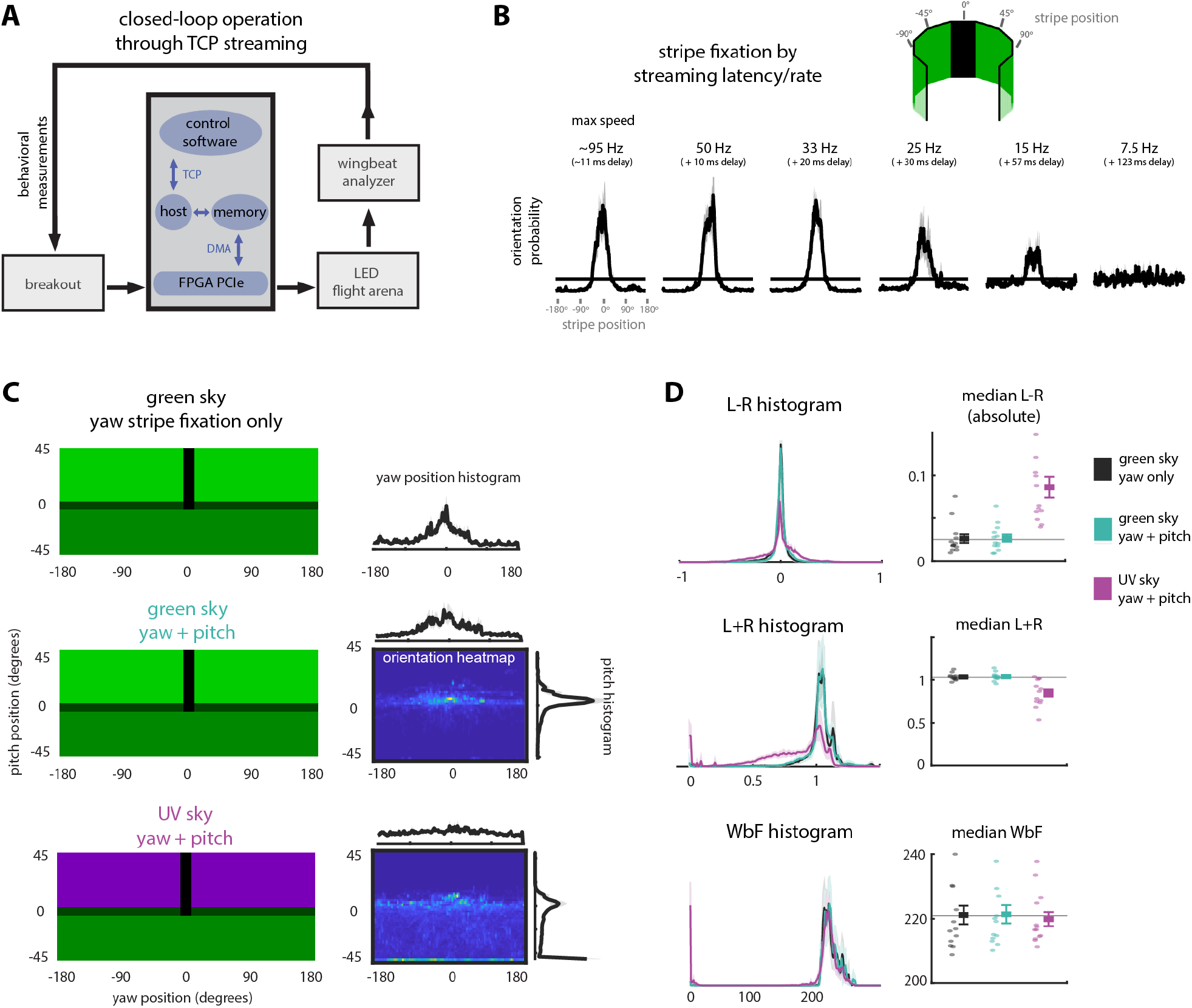
Virtual reality with flexible, interactive virtual environments. A) Schematic of the high-level control for closed-loop Virtual Reality (VR). In each cycle of the ‘streaming’ mode, the control software acquires behavioral measurements through a low-latency TCP connection from the FPGA PCIe through the host application, then renders a new pattern frame that is streamed over TCP to the host application, which sends the frame to the FPGA over DMA. The FPGA then sends the new frame to the panels (over SPI). This complete cycle can execute in <11 ms (for 1-bit patterns). B) Orientation behavior of flies flying in a closed-loop stripe fixation experiment rendered with the high-level VR control system, with various latencies (0, 10, 20, 30, 57, and 123 ms) added to reduce the closed-loop streaming update rate. Orientation histograms show the mean ± SEM heading probability for N = 11 flies. C) Three virtual environments composed of a bright sky, darker ground and horizon, and a single dark vertical stripe: environments 1 and 2 are entirely green in color, whereas environment 3 has a UV-colored sky. A flying fly’s turning behavior (L-R) controls the yaw rotational velocity of the environment in all 3 environments, while in environments 2 and 3 the fly’s pitch behavior (measured as L+R) controls the pitch angle of the environment, capped at ±45°. Single-dimension Pitch and yaw orientation histograms of flying flies in each of the 3 environments, and two-dimensional orientation heatmaps for environments 2 and 3. Histograms are generated from the combined data of N = 13 flies. D) Summary of behavior measurements (mean ± SEM of N = 13 flies) as histograms (left) and median values (right, with single fly values) for flying behaviors in each of the 3 environments shown in (C).

## Results

### The G4 modular display system

The G4 display system (Figure 1A) is composed of LED *panels* arranged in *columns*, that are grouped to form *arenas* of different shapes, such as flat walls or cylinders, that are quite compact and can therefore be integrated into a variety of experimental setups (Figure 1B).

Each panel is a ‘sandwich’ of a *driver* board housing the LEDs and a *communication* board that connects to other panels. G4 has ∼2.5 times the pixel density of the G3 LED modules^13^ (16×16 LEDs over 40×40 mm vs. 8×8 LEDs over 32×32 mm), yielding higher resolution visual arenas. Faster data transmission through SPI interfaces replaces I^2^C used in the prior G3 design. The G4 display is controlled by custom software and an FPGA that drives 12 parallel SPI busses, achieving sustained data rates per panel that are >20× faster than G3, supporting display refresh rates of 500 or 1000 Hz. By using surface-mount LEDs of widely available standard sizes, G4 panels can be customized with a variety and even combinations of LEDs.

A typical 9 column × 4 row, 36-panel G4 arena reliably operates at 500 Hz refresh rate while streaming visual patterns with 16 brightness levels and 1 kHz refresh rate for patterns containing 2 brightness levels. In our display system the refresh rate describes the number of unique frames shown per second (Figure 1C). Fine-scale control of brightness is achieved by modulating the LEDs’ duty cycle within a frame through the ‘stretch’ value (Figure 1C). The controller streams patterns directly to the arena, eliminating the need for intermediate buffers, and serves as a flexible data acquisition (DAQ) system, logging up to 8 analog inputs at user-specified sample rates (up to 200 kHz). Additionally, the controller implements up to 8 analog outputs that are precisely synchronized with display refresh (Figure 1D).

To allow researchers to rapidly use the G4 system without directly programming lower-level components, we developed graphical user interfaces and high-level software tools in MATLAB that are supported by extensive documentation and tutorials (https://reiserlab.github.io/Modular-LED-Display/G4). This open-source software provides users with interactive tools for generating visual stimuli, designing and running experiments, and processing the resulting data from within user-friendly, well-documented tools (Figure 2). Advanced users can extend these capabilities through scripts that provide further flexibility, such as modifying protocols mid-experiment, and direct access to the arena’s host application through low-level TCP/IP commands.

The G4 display’s combination of fine brightness control, fast refresh rates, and precise synchronization between the display and input/output channels, all controlled through high-level software, enables a range of experiments not previously possible. In each of the following sections, we introduce a biological question about *Drosophila* vision along with the technical challenges required to address it. We then describe our experiments that use unique features of the G4 display system to address these open questions.

### Determining the upper limits of *Drosophila* motion vision using high speed optic flow stimuli

The optomotor response of flies, in which the animals orient their head and/or their body to compensate for large-field visual motion, has been a critical behavioral tool for understanding the computational properties of insect motion vision^4,28,29^ and has recently been used to establish specific roles for specific visual neurons^30–33^. However, many substantial issues remain unresolved, such as whether the speed tuning of visual system neurons can explain the speed tuning of optomotor behavior^34,35^ and whether all axes of visual motion are identically speed tuned. In prior studies, the limited speed of the display prevented experiments testing the behavioral responses to high velocity motion^31,36^, and consequently the maximum stimulus speeds that flying flies can perceive are not well established.

Using the G4 display’s 1 kHz refresh rate, we presented grating stimuli moving over a wide range of speeds (>1000°/s) with corresponding temporal frequencies (the frequency of alternating bright/dark bars in a drifting grating) of up to 62.5 Hz (Figure 3A), which approaches the flicker fusion frequency of *Drosophila*^24^. Since the display’s refresh rate is independent of the pattern’s complexity (another improvement over G3), we examined responses to motion stimuli simulating rotational and translational motion around and along the 3 cardinal axes (6 flow fields, 2 directions each; Figure 3B). We presented these patterns in a randomized order to thorax-tethered, flying flies while recording the amplitude and frequency of the left and right wingbeats measured by an optical wingbeat analyzer^8^. The acquired behavioral data and display refresh cycles are precisely synchronized at 1 ms intervals. These open-loop trials were interleaved with closed-loop trials (see Methods and Movie 1) during which the fly’s turning controls the rotation of single bar typically positioned directly in front (called ‘stripe fixation’). These closed-loop trials serve to engage the fly and maintain vigorous flight while also conveniently resetting wingbeat amplitude differences close to zero, acting as an experimental strategy for standardizing the behavioral responses to subsequent visual stimuli.

In response to bilaterally asymmetric stimuli (yaw and roll rotations, lateral slip translation) flies mainly turn (quantified as the difference between Left and Right wingbeat amplitude, ‘L-R’ in Figure 3B,C) while responses to bilaterally symmetric stimuli (pitch rotation and lift translation) were primarily changes in the symmetric sum of wingbeat amplitudes (‘L+R’) as well as the wingbeat frequency. These results are largely in agreement with responses to impulsive visual motion along the 6 directions^37^, although noteworthy differences in the relative amplitude of the responses were found: the impulsive stimuli resulted in the largest turns to slip, and smaller yaw and roll responses, while our stimuli resulted in steady-state response levels (Figure 3A) that were similar for these 3 turn-inducing motion patterns (Figure 3B). These results clearly show that different visual motion patterns result in distinct responses and that flying flies steer by changing different combinations of wing stroke sum and difference (Figure 3C and Movie S1). Importantly, we have (for the first time) recorded the behavioral reactions of flying *Drosophila* to different visual motion stimuli over a speed range that includes stimuli faster than the flies’ ability to detection motion (Figure 3C). By comparing the speed tuning curves for the 5 (of 6) motion types that produced large behavioral reactions (thrust reactions were far more subtle; Figure S1) we find that temporal frequency tuning to different visual stimuli is rather consistent (Figure 3C, right), peaking at ∼8 Hz, and with a remarkably similar roll-off in response to higher motion speeds. These findings are largely in agreement with the peak tuning of the yaw and thrust behaviors of walking flies^38^, although we did not vary spatial frequency as was done in that study, and walking flies appear to be much more sensitive to thrust motion than we found in our flight measurements. The diversity of response levels to slower stimuli may reflect gain differences or contributions from visual pathways with distinct tuning. Nevertheless, the similar temporal frequency optimum for flight reactions to different patterns of visual motion, provides strong support for the view that a limited set of neurons (presumably the directionally selective T4 and T5 cells) set the tuning properties of diverse behavioral responses.

### Mapping the directional tuning of visual projection neurons using a motorized visual display

A common challenge to describing visual stimulus-evoked neural activity is determining how local stimulus features are integrated across visual space. The Local Preferred Direction (LPD) of Visual Projection Neurons (VPNs) can be mapped using a small moving stimulus at many points around the animal^12,39^. However, not all neurons are sufficiently driven by these hyper-local stimuli, and more complex combinations of input stimuli cannot be tested with this approach. We designed a motorized G4 display that overcomes these limitations by presenting larger, more flexible visual stimuli across the field of view, while working within the highly constrained space of a typical 2-photon imaging microscope. We applied this setup to characterize the motion response properties of a recently discovered VPN class^40,41^.

Our setup uses a rotatable, hemi-cylindrical (12 columns out of a circumference of 18, 20° per panel in azimuth) G4 arena. The entire display can be pitched up or down, while rotating around the position of the fly at the center (Figure 4A, top). The rotation of the arena is motorized for precise, repeatable, programmatic presentation of stimuli (see Movie 2). To stably access neurons in the head capsule, flies are typically mounted to a tapering fixture that occludes the dorsal field of view^42,43^. To take further advantage of the larger field of view of the motorized display, we designed a custom-machined conical head-mount (Figure 4A, bottom; design files shared at https://reiserlab.github.io/Component-Designs/physiology). This mount substantially improves the flies’ field of view while maintaining adequate access for the surgical preparation and subsequent placement of high-magnification objectives (Figure S2 and Movie S2). By presenting stimuli across the fly’s unoccluded field of view (Figure 4B,C), both in one location at a time and in multiple locations to assess global integration properties, we could more comprehensively map the directional tuning properties and receptive fields of VPNs.

Applying this technique to the recently described, regressive-motion-selective cell type LPC1^40,41^, we confirmed that regressive (i.e. back-to-front) motion presented to the eye ipsilateral to the imaged LPC1 glomerulus resulted in strong LPC1 calcium responses (Figure 4D,E top), while motion on the contralateral eye did not (Figure 4F, blue). Additionally, when a regressive (i.e. activating) motion stimulus on the ipsilateral eye was paired with 2^nd^ motion stimulus in a different location simultaneously, we found that progressive (i.e. front-to-back) motion on the contralateral eye consistently reduced LPC1 activation (4E, bottom). By mapping the activating and modulating motion directions in multiple visual field locations on the ipsilateral eye, contralateral eye, and binocular overlap region (Figure 4F, individual fly responses in Figure S3), we found the LPC1 population to be highly selective not just to regressive ipsilateral motion, but to the global pattern of optic flow produced by back-to-front translational motion (Figure 4G). This method, relying on a motorized G4 display and low-occlusion head mount, can be used to map the excitatory and inhibitory directional receptive fields of any visual projection neurons (or their targets in the central brain) over a wide field of view.

### Exploring the interactions between UV and green visual motion responses using precise brightness control

Commercial displays designed for humans are inadequate for studying color vision in animals that see wavelengths outside of the human visible spectrum, notably into UV. G4 display panels are designed for standard surface-mount LEDs (0603 package), so that panels can be assembled with different LED combinations. We built an arena using 2-color panels with a checkerboard arrangement of short-wavelength UV (365 nm) and green (570 nm) LEDs, to explore the relationship between color and motion vision in *Drosophila* (Figure 5A). This combination of LEDs is also well suited to mice and zebrafish spectral sensitivities^21–23^. Since LEDs provide uniform illumination for each color channel and from each panel, we arranged the panels into an experimentally convenient arenas (e.g. a hemi-cylinder) without the need to compensate for the significant differential scattering of short vs. long illumination wavelengths that complicate the implementation of projected displays^44^. Using our new 2-color display, we found that flies responded with robust open-loop optomotor turns and closed-loop object tracking (‘stripe fixation’) to visual patterns displayed on either the UV-only or green-only LEDs (Figure 5B), confirming and extending prior results^44–46^.

We tested the contribution of UV and green wavelengths to the perception of visual motion by testing whether they can be balanced at a point of isoluminance. At this condition, the relative motion of objects rendered with different wavelengths cannot be detected. Pixelwise intensity control makes the G4 system ideal for asking whether intensity combinations exist where the opposing motion of UV and green patterns exhibit isoluminance. We found that for bright green and dim UV combinations, flies turn in the direction of green. Similarly for bright UV and dim green they turn in the direction of the UV pattern motion (Figure 5C). However, we could not find an isoluminance point with these modest steps in relative intensity between the green and UV patterns tested. To see whether we could reliably establish isoluminant conditions, we used “stretch” modulation of a UV-only and green-only grating pattern displayed on alternating refresh cycles (halving the effective refresh rate of this 2 brightness level pattern to 500 Hz). While holding the green brightness constant, across trials, we adjusted the UV brightness in <1% increments. With this fine intensity control we found that UV/green isoluminance is indeed a reliable feature of the *D. melonagaster* optomotor visual response (Figure 5D).

### Virtual reality with flexible, interactive virtual environments

In the previous sections we demonstrated how G4 setups display visual stimuli to head-fixed animals with precision and high speed in open-loop (‘stimulus-response’) experiments. However, we also designed the system for low-latency closed-loop operations. Multiple closed-loop modes are implemented for customizable virtual reality (VR) experiments, where visual scenes of arbitrary complexity can be flexibly controlled to provide interactive visual feedback. A common VR experiment for flies uses rotational closed loop, in which a single behavioral measurement (flight: turning measured as ‘L-R’ Left–Right wingbeat amplitudes^47^; walking: rotation of air-supported ball^42,48^) is fed back to control the rotational velocity of the rendered virtual environment. Stripe fixation is a paradigmatic example, in which flies control the azimuthal position of a vertical bar, and preferentially orient their flight towards it^47^. This experiment is readily implemented in the G4 system, by pre-rendering all possible rotational positions of the stripe pattern and using the Left-Right signal to set the rotational velocity (scaled by a gain) that instantaneously sets the index into the ordered pattern frames^13^. To achieve low-latency closed-loop, this process is fully implemented by the FPGA controller which achieves a true input/output (I/O) latency of ≤2 ms (measured as the lag from analog input changes to changes in the display using a photodetector, all logged together). Additional closed-loop modes add a time-varying open loop term to the closed-loop analog input or use the analog input to directly set the pattern position (instead of setting the rate of position changes) with similarly low latency. The G4 system also supports a closed-loop ‘streaming’ mode where high-level control software (MATLAB in our implementation) renders new visual pattern frames in real-time based on behavioral measurements (up to 4 analog input channels, although other sources such as video-based postural measurements^49^ or spherical treadmill tracking data^50^ could be incorporated with further software development), while communicating with the FPGA controller’s host application over TCP/IP (Figure 6A). This streaming mode has much lower I/O latency (∼11 ms for 1-bit frames with 2 brightness levels and ∼16 ms for 4-bit frames with 16 brightness levels) than display systems using video graphics cards. This method gives experimenters full control over how the animal’s behavior determines what the animal sees.

How fast does closed-loop VR need to be? This fundamental technical specification can only be empirically assessed with a very fast feedback system. As a direct test of this requirement, we used G4’s streaming mode to assess flies’ stripe fixation with different feedback delays (for constant closed-loop gain). At the fastest closed-loop rate (95 Hz, ∼11 ms I/O latency) flies exhibited robust orientation to the dark bar (Figure 6B). Surprisingly, this behavior was unaffected by substantially increased I/O latency. Flies could reliably orient towards the bar when its position updated at 50 Hz or 33 Hz, with correspondingly longer delays (additional 10 ms and 20 ms) between the animal’s measured behavior and the display’s update. However, the orienting behavior noticeably degraded with 25 Hz updates, while only modest tracking was seen with 15 Hz updates, and no tracking at all was seen for 7.5 Hz updates (Figure 6B). For stripe fixation behavior in flying flies, we find that a closed-loop update of 33 Hz (∼30 ms I/O latency) or faster is compatible with stable object tracking, while longer delays apparently break the ‘realism’ of this virtual reality experiment. Flight behavior is notoriously fast, and we expect that more complex tasks, such as those defined not by the position of features but instead by their motion (such as optic flow), may require lower closed-loop latency. Even these requirements should be well within the capability of the G4 system, which we demonstrated is more than fast enough to accommodate VR experiments in *Drosophila*.

Using the G4’s ability to flexibly render scenes and record multiple channels of behavioral responses in near real-time, we set out to create more complex, naturalistic closed-loop experiments by extending the stripe fixation closed-loop task to 2-dimensions. The environment (generated using 16 brightness levels) was composed of a dark vertical stripe over a green background consisting of a bright sky, and a slightly darker ground, which are typical characteristics of sunlit open woodlands^51^. Tethered, flying flies controlled the azimuthal rotational velocity through L-R differences (Figure 6C, top). Building on this, we then allowed flies to control the pitch position of the pattern, allowing the horizon to drift up and down through changes to L+R, which we showed is related to pitch reactions (Figure 3B) and is known to contribute to body pitch control in flight^52^. Flies flying in this virtual environment controlled their position in both dimensions. They mainly oriented towards the stripe, while also holding a relatively constant pitch angle, with a preferred position for the horizon (Figure 6C, middle). We further enhanced this closed-loop simulation by replacing the green colored ‘sky’ with UV of a similar apparent brightness (informed by the isoluminance experiment, Figure 5). Under this UV-sky condition, stripe fixation was reduced and the pitch angle was much less stable, with flies often pitching fully downward (Figure 6C, bottom). Across all 3 conditions, the flies’ motor output was similar, except that the ‘UV sky’ condition resulted in strong reductions in flight vigor (measured by L+R, Figure 6D), suggesting perhaps that the sky is too bright. By pre-rendering the individual elements of these complex scenes (sky, stripe, horizon, and ground) and then arranging and coloring them in real-time according to the experimental condition and the animal’s estimated heading in the virtual environment, the frame rendering times are <1 ms, maintaining the system at near minimum total latency (∼17 ms), demonstrating the G4 system’s flexibility in powering these elaborate, interactive VR experiments.

## Discussion

Historically, vision scientists have used different display strategies for distinct purposes. In this study we demonstrate a new, high-speed, modular display system, that can be readily adapted to the full range of laboratory experiments for studying vision in small animal models (Figure 1,2). We demonstrate the unique capabilities of this system while exploring a broad set of questions in *Drosophila* vision: (1) using fast display rates we report the complete speed tuning of flight reactions to different patterns of visual motion (Figure 3), (2) with a panoramic, motorized display we map the spatial response properties of visual projection neurons across the fly’s field of view (Figure 4), (3) using precise intensity control and a UV-green variant of the display, we show that isoluminance for these wavelengths is a reliable feature of *Drosophila* vision (Figure 5), and finally (4) using low-latency pattern streaming to implement several virtual reality assays, we demonstrate that our system’s performance is well in excess of the minimum required for high-fidelity closed-loop behavior in flies and further show the flexibility of the system to incorporate multiple behavioral measurements into the controlled simulation of free behavior (Figure 6).

Small, flat, reliable display systems are favored for fine-grained receptive field mapping in retinal or cortical circuits, while computer-graphics based systems, often accompanied by substantial input-output delays, are preferred for more panoramic virtual reality displays. Here we demonstrate a single system which is a general solution for all visual display needs in our laboratory, spanning receptive field mapping to detailed psychophysical tasks to closed-loop virtual reality simulations of free behavior. The technical improvements and advanced features of the G4 display system enabled new and improved experiments on visual information processing in *Drosophila*. Furthermore, the flexibility of customizing the displayed wavelengths opens future studies into color vision for many species and can be used to manage complex wavelength requirements of modern experimental setups integrating optogenetic with visual stimulation together with different fluorescent indicators. This new system takes important steps toward the goal of bringing more complex and naturalistic visual stimuli into the laboratory without sacrificing control or precision.

## Acknowledgements

We thank Igor Negrashov and Bill Biddle for help in designing and fabricating the conical head mount, and Jon Arnold for designing the motorized rotational stage. We thank Aljsoscha Nern for the LPC1 split-GAL4 line and Edward Rogers for help with fly husbandry. We thank Ofer Mazor and Pavel Gorelik for feedback and updates to the documentation and code for the project. We are grateful to members of the Reiser lab, especially Eyal Gruntman, for feedback on the design of the system and discussions on the manuscript. This project was supported by the Howard Hughes Medical Institute. This article is subject to HHMI’s Open Access to Publications policy. HHMI lab heads have previously granted a nonexclusive CC BY 4.0 license to the public and a sublicensable license to HHMI in their research articles. Pursuant to those licenses, the author-accepted manuscript of this article can be made freely available under a CC BY 4.0 license immediately upon publication.

## Methods

User documentation and technical resources for several generations of modular LED systems is organized at the website https://reiserlab.github.io/Modular-LED-Display/. Design files, bill of materials, and other details about the custom hardware and software for the G4 system are organized in the “Generation 4” menu entry at https://reiserlab.github.io/Modular-LED-Display/G4.

### Display system hardware

G4 Panels are composed of two parts: a *driver* board containing a 16×16 square grid array of LEDs, and a communication or *comm* board which receives pattern data from a controller.

The *driver* board with its 16×16 surface mounted LEDs (0603 form factor) is internally subdivided into four 8×8 LED quadrants. For each quadrant of LEDs one ATMEL ATmega328 microcontroller unit (MCU) delivers the row and column patterns to drive the quadrant of LEDs. The electronic design of each quadrant of the driver board is based on the G3 panel LED driver design, where one ATmega also drives 8×8 LEDs^13^.

With a second PCB in each panel, the G4 design improves the communication with the controller, which was a bottleneck in G3 refresh rates. The *comm* board houses another ATMEL ATmega328 and serves as a high-speed communication interface, receiving pattern data formatted for simple partitioning through an SPI connection, a common communication standard typically set to operate at 4 MHz in our setup. The data for each driver quadrant is sent via SPI (also operating at 4 MHz) to the four MCUs on the driver board, well within the 1 ms target refresh cycle per panel. Up to eight comm boards can be daisy-chained together to form a “*column*.*”* Whereas in the G3 design each panel needed to be addressed over the I^2^C bus, the G4 system panels are automatically addressed through their location in the column. Within each column eight different chip select lines are passed through the comm boards, shifting by one position between each row to make a distinct line the active chip select. To create a full-size display, up to 12 columns are connected to a larger “arena board” where each column has its own dedicated SPI signal.

### Low-level PC control of display system

The up to 12 individual SPI busses that drive the columns of an arena, are supplied via a direct connection to the digital output bus of a reconfigurable I/O module packaged as an FPGA PCIe card (National Instruments PCIe-7842R). This device can deliver data in parallel to all 12 channels at above the required bandwidth of 4 MHz, while maintaining clock and signal synchronization. The PCIe card is in a host PC (tested on Microsoft Windows 7 and 10) that allows for flexible high-level software development and control. We prototyped the display controller for the integrated system using other microcontrollers, but turned to a single FPGA-based reconfigurable I/O module for synchronized control of 12 high-speed busses.

There are two custom software components developed in LABVIEW (that are distributed as a binary package) – firmware that is deployed to the FPGA, and a host application (called “HHMI G4 Host”) that runs on the Windows PC and manages all communication with the FPGA. The core function of the FPGA is to stream pattern data at frame rates of 1 kHz (1-bit, 2 brightness levels) or 500 Hz (4-bit, 16 brightness levels) via SPI to each of the 12 columns. These frame rates are set by the FPGA firmware, and with further optimizations could be increased if necessary. The reconfigurable I/O module also houses data acquisition (DAQ) hardware that supports up to 8 analog input and 8 analog output channels with up to 200 kHz sampling rate. In our typical use of the G4 controller we log up to 4 analog input channels (−10 V to +10 V) at sample rates up to 20 kHz and generate up to 4 16-bit analog output channels that are streamed out at the display refresh rate. The analog channels are logged together with the timestamps for each display refresh cycles with 1 µs precision.

Pattern data stored on the host PC is sent to the FPGA PCIe card via Direct Memory Access (DMA) and “directly” routed to the FPGA’s implementation of 12 SPI channels. With this design, and optimized buffer management, a full frame of data is sent within each 1 ms update cycle. The PCIe card’s firmware is entirely controlled through the host application, which is accessible through a TCP/IP protocol. The different TCP/IP commands specify what frame or sequence of frames (each referred to as “*patterns*”) to display and in what “*Display Mode*.” Display modes determine the logic by which the order and timing of pattern frames are shown on the display, either in open-loop or closed-loop, as described in Table 1 (more in-depth technical descriptions of each mode are available https://reiserlab.github.io/Modular-LED-Display/G4/Display-Modes). In open-loop modes, patterns are displayed in a predetermined way, such as displaying a single static frame for a set duration, cycling through multiple frames at a constant rate, or by presenting frames in a pre-set order stored as a “position function” (see “*Display system software*” section). In closed-loop modes, an external signal acquired as an analog input determines which frames of a pattern are presented (after some scaling) in the following display updates (see Figures 5 and 6 for an example of closed-loop landmark fixation).

**Table 1:** G4 Display Modes. List of display modes of operation used by the G4 system, with basic descriptions of how each mode controls how pattern frames are displayed. The rows of open-loop (i.e., predetermined) modes are shown in white, while closed-loop (i.e., modulated by analog input) modes are shown in gray.

|  |  |
| --- | --- |
| Position function | Uses a position function (array of frame indices) to determine which frame number of a pattern will be displayed at each refresh cycle of the display |
| Constant frame rate | Cycles through the pattern in order at a constant frame rate |
| Static frame | Shows only a single frame from the pattern |
| Closed-loop sets frame rate | Cycles through pattern frames at a rate set by the ADC0 analog input voltage (modified by a gain and offset), with negative values cycling from higher to lower frame indices |
| Closed-loop + position function | Combines “closed-loop sets frame rate” with pattern frame position set by a position function |
| Closed-loop sets frame index | Displays pattern frames by scaling the analog input voltage to the pattern frame indices |
| Closed-loop streaming | Displays frames streamed from the high-level control software over the TCP/IP connection as fast as new frames can be delivered, up to 100 Hz, while sending 4 analog inputs back at the same rate to be used by the control software to generate new frames |

The host application listens for TCP/IP commands, which are flexible and allow the use of alternative high-level platforms for communicating with the host application. The host application loads the firmware to the FPGA on the I/O card, configures the firmware for specific display modes, and is used to start and stop the different conditions after which the FPGA firmware runs autonomously with direct access to the computer storage via DMA. The closed-loop streaming mode is an exception, where frames are received via TCP/IP and passed on to the FPGA. The host application also provides a simple GUI to interact with the system which internally sends the same TCP/IP commands specified in the protocol. In the next section we describe a MATLAB-based user-friendly high-level method to interact with the host application.

### High-level display system software

For increased user-friendliness, the TCP/IP protocol commands of the host application, that ultimately controls the FPGA and SPI commands sent to the panels, are accessible via high-level MATLAB control software. MATLAB can be used to design, render, and save patterns prior to an experiment and to handle the logic of the experimental structure. In addition, MATLAB can access analog input data during the experiment (down-sampled to 100 Hz), for example to plot data during an experiment or to create custom closed-loop streaming experiments (see Figure 6).

The software tools developed for the G4 display system are organized to follow the typical process of creating visual patterns (“G4 Pattern Generator”), defining their temporal arrangement (“G4 Function Generator”), defining an experimental protocol (“G4 Protocol Designer”), running experiments (“G4 Experiment Conductor), and processing logged data (Figure 2). We note that these tools can also be used independently to simplify specific tasks in cases where an advanced user wishes to further customize their experiments.

The “G4 Pattern Generator” determines what a pattern (either a single-frame or multi-frame sequence) will look like, based on user-defined parameters. This GUI tool allows for rapid visualization and generation of various commonly used patterns such as lines, edges, gratings, and star fields, parameterized for example by size, brightness, and movement direction. Using a 3D customizable coordinate-based pixel map of the arena, patterns can be generated that simulate rotation around or translation along a fixed point within the arena. The resulting patterns are presented either on the entire display (e.g. see Figure 3) or in restricted spatial locations within the simulated visual field (e.g. see Figure 4). To extend the functionality of the G4 Pattern Generator, users can modify, combine, or create new patterns from scratch using scripts.

The “G4 Function Generator” can be used to specify when and how long each frame of a pattern is shown on the display. These “Position Functions” set the temporal “position” of each pattern frame relative to the display’s refresh rate; when using 16 brightness level patterns at 500 Hz refresh rate, frame positions will be set for every 2 ms, while patterns with two brightness levels at 1 kHz refresh rate will be set for every 1 ms. Position Functions determine the order that frames of a pattern are displayed, as well as the parameters defining pattern motion (e.g. constant speed drifts or sinusoidal motion). Additionally, the “G4 Function Generator” tool can generate Analog Output (AO) Functions: just as Position Functions control the displayed frame for every refresh cycle, AO functions control the voltages of 4 (extendible to 8) analog output channels for every refresh cycle, allowing precisely synchronized external equipment.

The “G4 Protocol Designer” is used to define the order in which the previously generated patterns are sent to the display as part of an experiment protocol. A “G4 protocol” can be as simple as an ordered list of patterns to be streamed to the display. A more sophisticated protocol may contain associated Position or AO Functions, queueing patterns in randomized repeating blocks, with different display modes, separated by delay periods, custom inter-trial patterns, and closed-loop trials. Each trial can be previewed to verify that the selected pattern is correctly presented, either in a simulated display on the PC screen or on the G4 arena. When saving an experiment, the “G4 Protocol Designer” generates the information and folder structure accessible by the host application and FPGA (via DMA) that are necessary to run the experiment. The actual experiments can be initiated through another GUI application, the “G4 Experiment Conductor”.

The “G4 Experiment Conductor” is a GUI to run experiments. It asks the user for additional metadata about the current instance of the experiment such as the experimenter’s username, timestamp, and details of the experimental animal. After starting the experiment, a progress bar provides feedback. The Conductor can also process and plot streamed analog input data once individual trials are completed. This provides additional quality control mechanism by visualizing the collected data as it is being logged.

When an experiment has completed, a set of acquired data, including timestamped frame positions and analog input channel data are logged as .tdms files (Technical Data Management Streaming, a National Instruments file format). We provide additional data processing MATLAB scripts to load these files into the MATLAB workspace, convert the data to the .mat file format, and plot basic analysis in several standard ways (e.g., time series, histograms, and trial averages). If the desired plot formats are already known during the protocol design, the parameters for processing and plotting data can be pre-determined from within the G4 Protocol Designer tool and executed immediately after an experiment has completed. This standardization greatly simplifies organizing experiments across setups and experimenters.

### System Validation

Operation and performance of the G4 system was assessed using our typically sized 9 column, 4 row cylindrical arena, using standard rotational grating and single-stripe patterns (e.g. Figure 5B). Display brightness and LED spectra (Figures 1C, 5A) were measured using a spectrometer (Ocean Optics USB4000) with a fiber optic cable positioned over a single LED in the center of the cylindrical arena. The timing of display refresh cycles and analog outputs to the internally logged pattern positions (Figure 1D) was calculated using the measurements of a fast photodetector (Thorlabs PDA10) logged as an analog input channel, with an analog output wired back into the G4 system on a second analog input channel. Closed-loop latency was measured using an external function generator to deliver TTL pulses into a G4 analog input to switch pattern frames, where the delay between logged input and photodetector changes was measured.

### Experimental Animals

Flies were reared under standard conditions: 25° C, 60% humidity, 16h light / 8h dark, cornmeal agar diet. Behavior experiments were performed on female flies 3-5 days post-eclosion, while imaging experiments were performed on female flies 1-4 days post-eclosion. For behavioral experiments (Figures 3, 5, and 6), Dickinson Lab (DL) flies were used. The original laboratory culture was maintained in Michael Dickinson’s lab, from which the Reiser lab established a copy at Janelia in 2007. This strain has been used in dozens of behavioral studies and has been referred to as ‘DL’^14^. For calcium imaging experiments (Figure 4), flies expressing GCaMP6m in LPC1 cells were used^40^. Cell type specific expression was achieved using the Split-GAL4/UAS expression control system^53^.

### Preparation for tethered flight

Flies were prepared for flying behavior experiments as previously detailed^13,54^ and summarized here: Cold-anesthetized flies were glued to a tungsten rod (catalog #71600; A-M Systems) by the thorax with UV curing glue (KOA 300-1; Kemxert, Poly-Lite, York, PA, USA) and were given at least 30 minutes of recovery time at room temperature and brightness before testing. All experiments were completed within 5 hours of tethering.

### Preparation for in vivo calcium imaging

Flies were prepared for 2-photon imaging experiments similar to previously described experiments^42^ with some modifications as summarized here: Cold-anesthetized flies were tethered to a fine tungsten wire using UV-curing glue. The two most anterior legs (T1) were severed and sealed with glue to prevent the fly from grooming and obstructing its visual field. The tethered fly was positioned up to the opening of a custom-machined PEEK plastic conical mount to allow access to the back of the head for dissection and imaging while obstructing as little of the fly’s visual field as possible (Figure 4, S2; for CAD files see https://reiserlab.github.io/Component-Designs/physiology). The fly was glued to the open tip of the conical mount at the rim of the head and the upper thorax, sealing most of the opening. The proboscis was glued to the severed legs to further immobilize the head. The back surface of the fly’s head was bathed with fly saline (103mM NaCl, 3mM KCl, 1.5mM CaCl2, 4mM MgCl2, 26mM NaHCO3, 1mM NaH2PO4, 8mM trehalose, 10mM glucose, 5mM TES) and a hole in the cuticle was cut to expose the PLP region of the brain using a fine tungsten needle and holder (Fine Science Tools #10130-05, # 26018-17.) Muscles 1 and 16^55^ were severed to reduce motion of the brain within the head capsule and excess fat was removed from the surface of the brain.

### Visual Stimuli

Visual stimuli were presented to tethered flies using two configurations of the G4 display system. Behavioral experiments used an arena with the shape of ¾ of a cylinder, made of 9 panel columns (∼150 mm interior diameter), spanning 270° azimuth, while imaging experiments used a 2/3 of a cylinder arena, made of 12 panel columns (∼230 mm interior diameter) spanning 240° azimuth. For imaging experiments, to further separate the LED emission spectra from GCaMP6m fluorescence, a low-pass gel filter (Deep Golden Amber gel filter, LEE Filters #L135S) was laid on top of the LEDs along with a diffusing screen to prevent reflections from the filter. All patterns that were displayed on the cylindrical LED arenas were generated using the G4 Pattern Generator tools for parameterizing and visualizing moving grating patterns for this display system. Displaying these patterns at multiple different speeds/temporal frequencies was accomplished by changing the rate in which the frames of the patterns are cycled through, defined by Position Functions.

All grating stimuli consisted of square-wave gratings set to 60° spatial wavelength and 100% contrast – other parameters of the grating patterns (e.g. temporal frequency and color) are as described in the text. For tethered flight optomotor experiments (Figure 3 and 5B), a single trial of open-loop grating stimuli began with a 2 second pre-trial of closed-loop condition of a dark vertical bar (30° width) on a green background, called “stripe fixation”. This was followed by a 2 second trial of the moving grating stimulus, followed by another 2 second post-trial stripe fixation. The stripe fixation pre- and post-trial segments were interspersed between all moving grating trials to maintain robust flying behavior for the duration of the experiment. For UV-green competing grating stimuli, the closed-loop stripe fixation inter-trial only used the green LEDs for the stripe pattern. For calcium imaging experiments (Figure 4), a single trial consisted of a 2 second pre-trial of a uniform medium brightness frame, followed by a 1 second trial of moving gratings, followed by another 2 second post-trial of the uniform frame, after which the arena was turned dark for 4 seconds to allow the arena to pitch to its new position. The increased time in between trials in calcium imaging experiments relative to behavioral experiments allowed more time for the GCaMP6m fluorescence signal to return to baseline fluorescence levels. For stripe fixation and VR behavioral experiments (Figure 5B and 6), each trial consisted of 15 seconds of closed-loop stripe fixation beginning with the stripe in a random orientation around the fly, after which a 5 second inter-trial of stripe fixation took place at default pattern settings (green background), beginning with the stripe directly in front of the fly. All trials of visual stimuli to be presented to a fly were pre-generated and their order of presentation was randomly permuted, and every trial was repeated 3 times in a new permuted order (random block trial structure), after which a mean response for each experimental condition was calculated for each fly individually by averaging the 3 repetitions. These experiments were implemented with custom scripts, and these protocols formed the basis for the standardized functionality that is now supported by the new software tools described in the section “***High-level display system software*.*”***

### Pitching Arena

A (2/3) cylindrical arena with 230 mm interior diameter and panels in 12 columns covering 240° azimuth and 3 rows was attached on one side to a motorized rotary stage (Zaber X-RSW-E, Zaber Technologies) using a custom mount to pitch the arena around a tethered fly (Figure 4). The rotary stage mount was aligned to the center of the column 2, row 2 LED panel, while a free-rotating mount was aligned to the center of the column 11, row 2 LED panel, to create a stable axis of rotation through the center of the arena (mounts at ±90° to the sides of the fly). Rotation commands were sent to the rotary stage using a MATLAB plugin, allowing the arena pitch and G4 display system to be controlled within the same script.

### Tethered flying equipment and analysis

The methods used for behavioral experiments have been detailed previously^8,13^ but are summarized here. Tethered flies were positioned in a hovering posture (60° body angle) in the center of the LED arena. Wing motion was measured using a “wingbeat analyzer” that measures the amplitude of the left and right wingbeats using an optical detector of the shadow size created by infrared LED placed above the fly. The relevant behavioral outputs of the wingbeat analyzer are the instantaneous measurements of the wingbeat amplitude of the fly’s left and right wings, the instantaneous difference between the left and right wingbeat amplitudes (left minus right, “L-R”), and the frequency of wingbeats. These analog signals were acquired and logged at 1 kHz using the on-board data logging of the FPGA based reconfigurable I/O PCIe card used as the controller of the G4 display system (see section “Low-level PC control of display system”). The L-R signal was processed separately either using the on-board fast (1 kHz frame rate) closed-loop mode of the G4 system (Figures 3, 5) or in MATLAB for closed-loop streaming experiments (Figure 6). All flies were positioned such that the wingbeats as measured by the wingbeat analyzer and visualized through an oscilloscope were of similar shape and amplitude, and that the flies were able to fixate a stripe in closed-loop operation, where the L-R signal (a reliable measurement of yaw rotational torque) controlled the apparent rotation of a dark vertical stripe (30° wide) on a bright arena background.

Using the logged left and right wingbeat amplitude data, a time-series of “L-R” was calculated by subtracting the right wingbeat amplitudes from the left for every timepoint, and a time-series of “L+R” was calculated by adding the left and right. The time-series of L-R, L+R, and wingbeat frequency (WBF) measurements in response to multiple repetitions of each visual stimulus were averaged together to produce a mean response for each fly, to each visual stimulus. For the mean time-series of L-R, all flies were then normalized to the 98^th^ percentile, a robust estimator of the max, L-R measurement across all conditions, which scaled the values to range from approximately -1 to +1, where -1 is a near maximum leftward (or counter-clockwise) yaw turn and +1 is a near maximum rightward / clockwise yaw turn. For the mean L+R (and WBF) time-series, all flies were normalized to the mean L+R (WBF) measurement during the 2 seconds prior to the open-loop experimental visual stimuli (2 seconds of closed-loop stripe fixation immediately preceding the experimental stimuli), scaling the values from approximately 0 to +1, where 1 represents the baseline L+R (WBF) levels prior to the start of experimental visual stimuli. For the summary tuning curve plots, a single value of L-R, L+R, and WBF responses was taken from each time-series by averaging the mean time-series values during the 2 seconds of experimental visual stimuli. Both time-series and summary data are plotted as mean of all flies ± SEM. In some cases (Figure 3C, Figure 5B,C,D), a further averaging step is taken when bilaterally symmetric conditions, e.g. clockwise and counter-clockwise versions of the same stimulus condition, are averaged, but with one set of behavioral data also sign inverted (to preserve the turning direction).

### 2-photon imaging and analysis

LPC1 neurons in the posterior lateral protocerebrum (PLP) of *Drosophila* (Figure 4) were imaged using a two-photon microscope (Prairie Ultima) with near-infrared excitation (930 nm, Coherent Chameleon Ultra II) and a 40x objective (Nikon CFI APO 60XW). The excitation power varied between 18-22 mW at the sample. T-series of z-stacks (z-series) were taken at 128×128×5 pixel resolution (0.446 µm/pixel along x and y axes; 5 µm/pixel in z-axis) at a rate of 2.02 Hz; x and y dimensions were scanned with galvo-galvo scanning and the z-dimension with piezoelectric scanning. For each experimental condition, the acquisition of 10 z-stacks were triggered to start precisely 2 seconds before the start of the visual stimulus using the on-board analog outputs of the display system: 4 z-stacks were recorded in the 2 seconds prior to the experimental stimulus, 2 recorded during the 1 second stimulus, followed by another 4 after the stimulus had ended.

Z-series data was converted into a t-series by taking a mean z-projection of each z-stack. Forming a t-series by collapsing z-stacks minimizes the effects of brain motion in the z-direction. Motion in the x and y directions was corrected using “imregister” in the MATLAB image processing toolbox: each image was first gaussian filtered (alpha=3 pixels) to improve registration of noisy images and registered using the ‘multimodal’ metric and ‘rigid’ transformation. Using this motion-stabilized t-series, an ROI was manually selected from the entire LPC1 glomerulus, excluding any additional axon tract that didn’t overlap with the glomerulus. The mean pixel value within the entire ROI was used as the population average fluorescence for each frame. Since each experimental condition resulted in 10 values for each experimental condition (4 values before the stimulus, 2 during, and 4 after), the fluorescence change (ΔF/F) caused by a visual stimulus was calculated by dividing all 10 fluorescence values by the baseline fluorescence – the mean of the first 4 ‘pre-stimulus’ values. Peak ΔF/F was computed by finding the max value of values 5-10 in each time-series; values 1-4 are ignored as they are the pre-stimulus fluorescence change values. Both ΔF/F time-series and peak ΔF/F are plotted as mean of all flies ± SEM (Figure 4E). The average response to each direction and the effect of modulation by stimulation at a second location are converted to polar coordinates and the simple vector sum is used to summarize these responses (Figure 4F).

---

**Figure S1:**
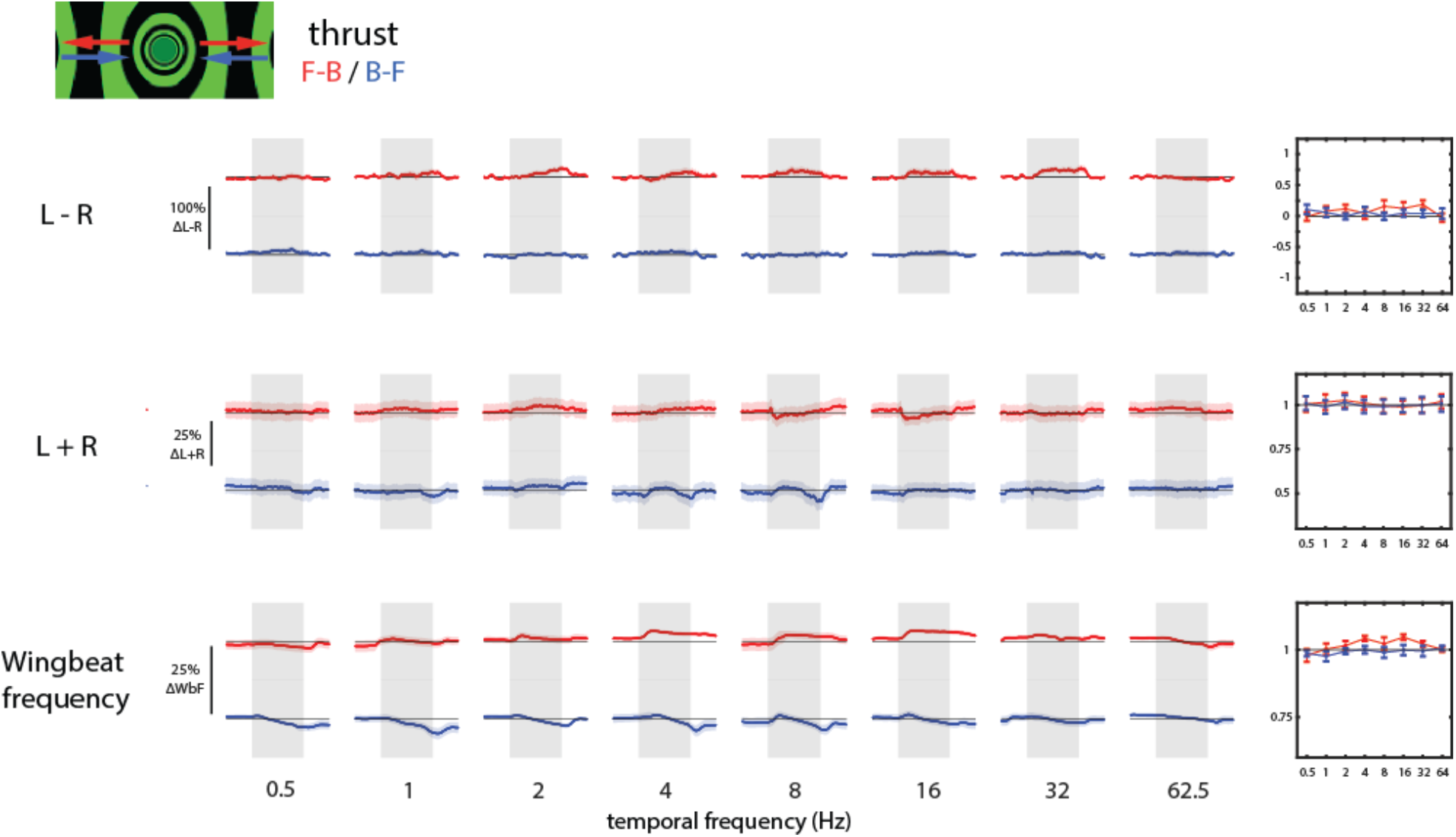
Small flying responses to thrust stimuli, related to Figure 3. Measurements of the left/right difference in wingbeat amplitudes (L-R), the left/right sum of wingbeat amplitudes (L+R), and the wingbeat frequency (WBF) in response to front-to-back and back-to-front drifting gratings (“thrust” motion) at 8 temporal frequencies. Each trial began with 2 seconds of stripe fixation (data not shown, see Methods for details), followed by 2 seconds of the specified drifting grating (gray shaded region), followed by another 2 seconds of stripe fixation. Responses are shown as mean ± SEM of N = 21 flies, with reactions to front-to-back thrust in red and back-to-front thrust in blue. Temporal frequency tuning curve summaries (mean ± SEM during the 2-second stimulus window, of N = 21 flies) of the behavioral reactions.

**Figure S2:**
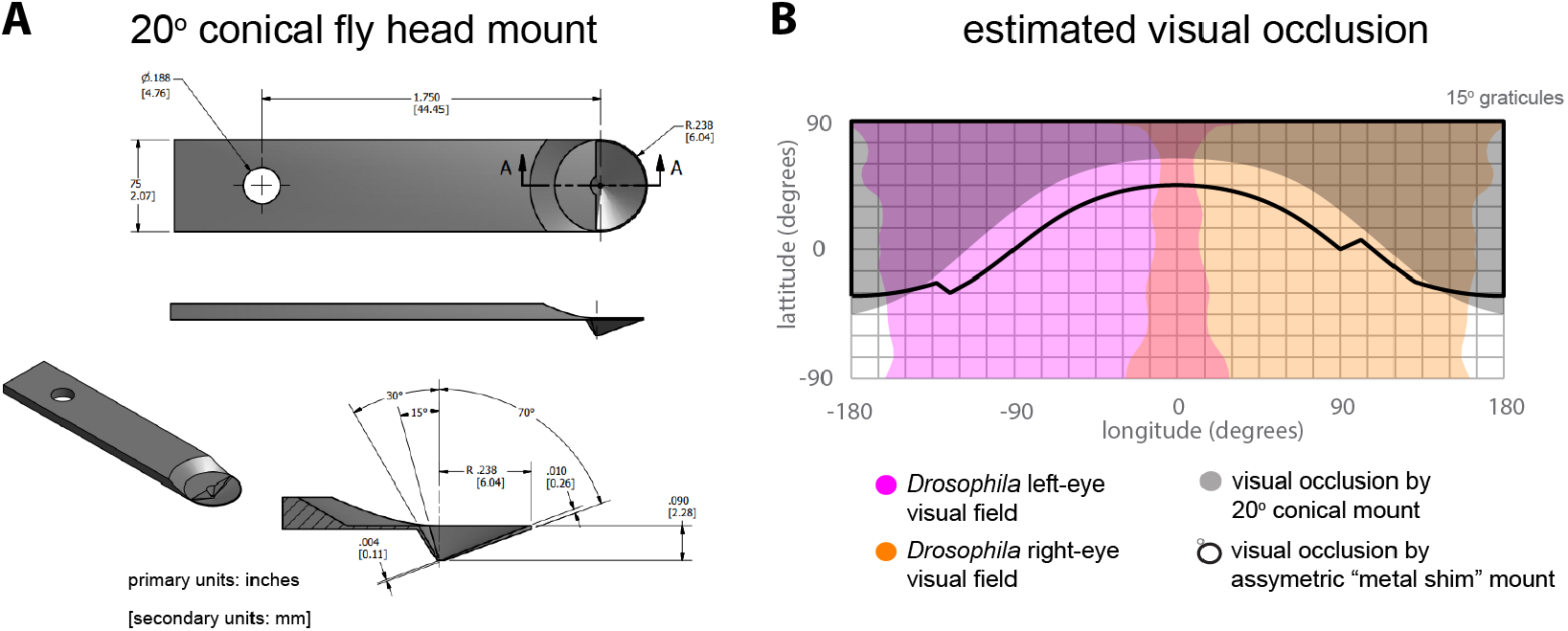
Conical fly head mount for decreased visual field occlusion, related to Figure 4. A) 3D rendered design schematics of the conical mount used for head-fixed calcium imaging experiments (Figure 4), shown from various angles. Designed to allow surgical and optical access to the head of a fruit fly from the dorsal side. B) Estimated regions of the *Drosophila* visual field which is blocked by the new conical head mount in (A), as compared to a previously used fly head mount built using a bent and laser-cut metal shim^42,57,58^. The conical head mount increases the field of view that can be visually stimulated and is left-right symmetric.

**Figure S3:**
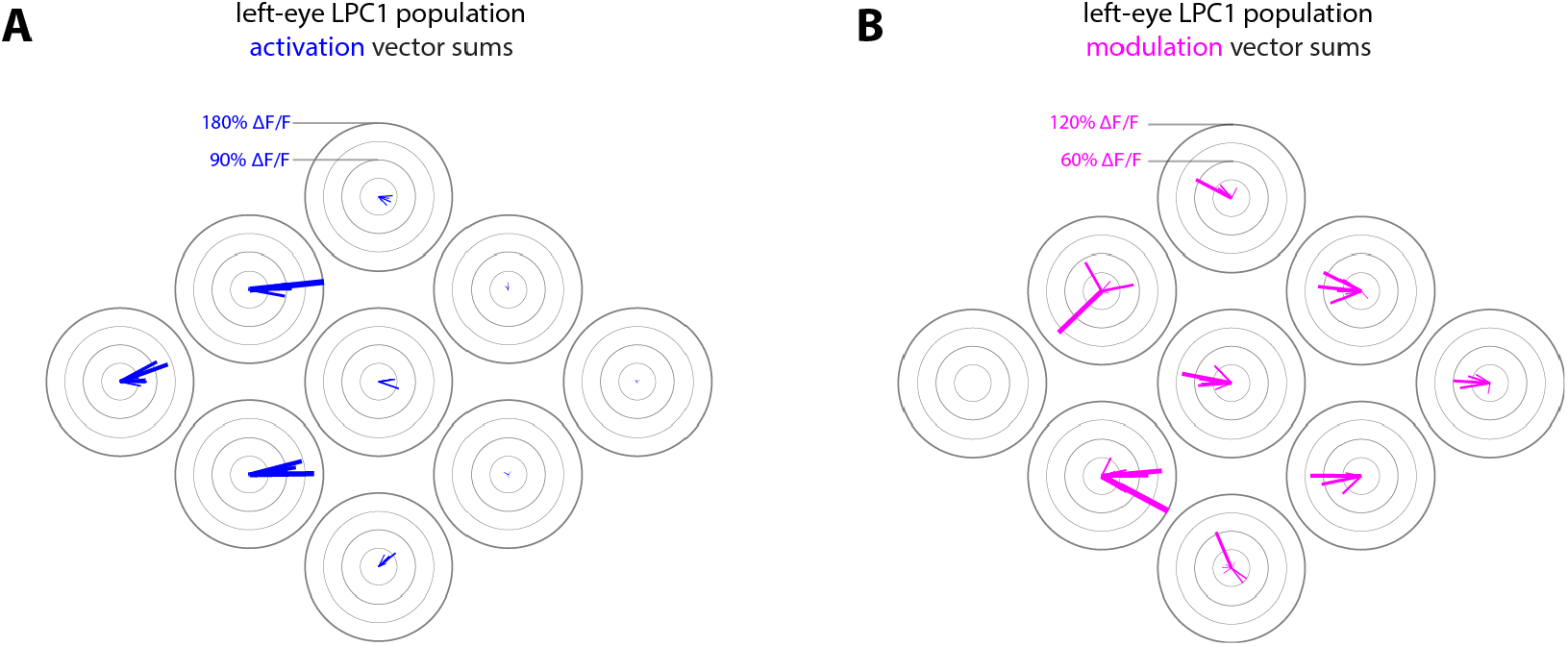
Individual LPC1 activation and modulation motion direction tuning maps, related to Figure 4. A) Vector sums of the left LPC1 response to 8 directions of motion stimuli presented in the 9 spatial locations shown in Figure 4B,C, plotted for each fly individually (N=8). The length of each blue line represents the response amplitude of the vector sum, while the direction of each line represents the motion angle of the vector sum. B) Similar to (A), vector sums of the left LPC1 modulated response resulting from simultaneous presentation of rightward motion in location E1 (producing a baseline level of activation) together with presentation of one of 8 directions of motion in one of locations E2-3, N1-3, and S1-3 (summarizing 64 experiments total – 8 directions at 8 locations). Modulation in E1 could not be calculated as E1 stimulation was reserved for baseline activation.

**Figure S4:**
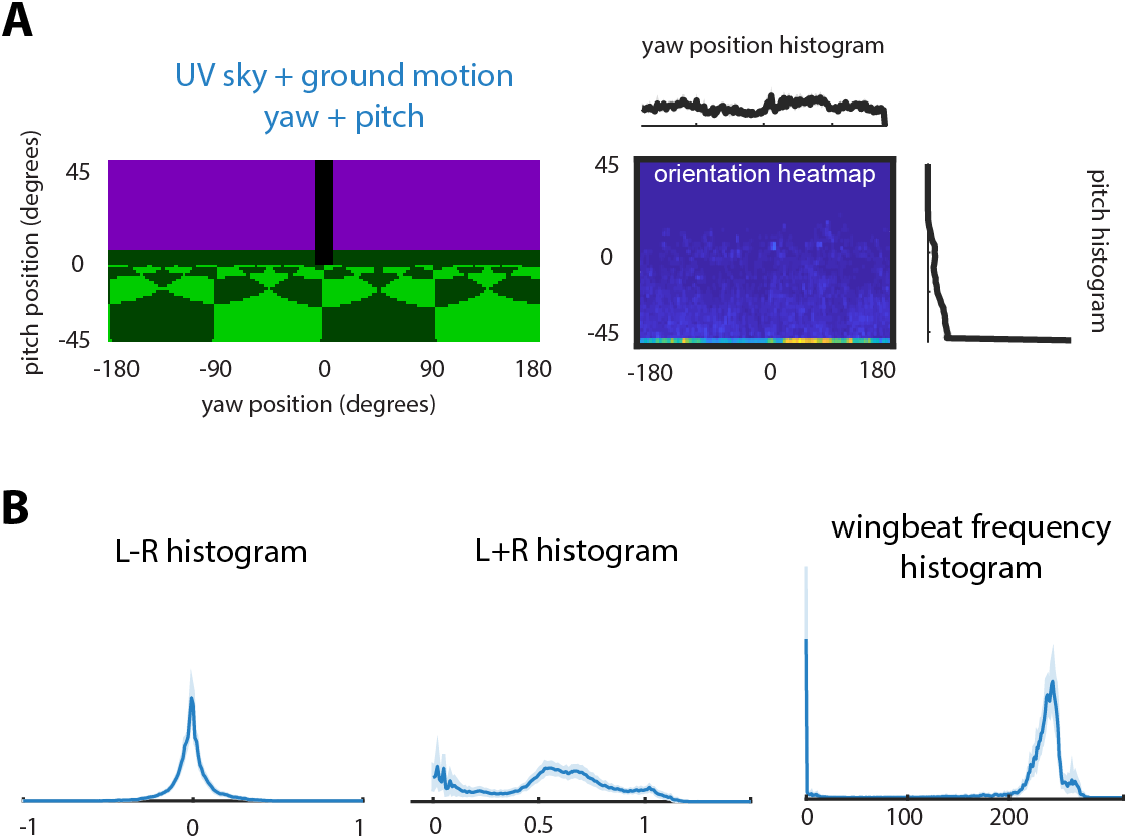
VR environment with UV sky and simulated ground-motion, related to Figure 6. A) A virtual environment composed of a bright, UV-colored sky, dark green horizon, checkerboard-textured green ground, and a single black vertical stripe. The textured ground is continuously shifting front-to-back to simulate the tethered flying fly’s forward translation. The fly’s turning behavior (L-R) controls the yaw rotational velocity of the environment while its pitch behavior (measured as L+R) controls the pitch angle of the environment, capped at ±45°. 2D (pitch and yaw) orientation histograms of flying flies in this environment, with separate yaw and pitch histograms shown above and to the right, respectively. Data is combined from N = 13 flies, completed as part of the same series as in Figure 6. Unlike in the experiments in Figure 6, in this condition flies did not orient towards the bar and persistently pitched downwards. B) Histograms of the 3 behavioral measurements (L-R, L+R, and wingbeat frequency) of flying flies (mean ± SEM of N = 13) in the environment shown in (A).

**Movie S1: Tethered flight behaviors in a cylindrical G4 arena, related to Figure 3**.

A tethered, flying fly responding to open-loop yaw rotation and lift translation stimuli, interspersed with closed-loop stripe-fixation trials. This movie is played back at 50% speed and is composed of two different synchronized videos combined to show the display arena as well as the flies’ reactions. The fly is suspended below an infrared LED that casts a shadow onto a pair of detectors below. These shadows are measured to quantify the amplitude of each wing’s stroke. Flies respond to right (clockwise) and left (counter-clockwise) yaw rotational motion stimuli by differentially modulating their wingbeat amplitudes (L-R), with little effect on the sum of left and right wingbeat amplitudes (L+R). Conversely, flies respond with large changes in L+R to up and down lift translation motion stimuli, with minimal L-R changes. During closed-loop stripe fixation, the actively orients towards the dark vertical stripe by turning. The L-R signal controls the rotational velocity of the stripe.

**Movie S2: Motorized pitching arena for adjustable visual field stimulation locations, related to Figure 4**.

A 12-column, 3-row G4 arena (240°, or 2/3 cylinder) connected to a custom motorized pitching mount, is adjusted to 3 different pitch positions. The pitch positions relate to the equatorial, north, and south positions relative to a mounted fly (head pitched downward to 45°) as shown in figures 4A,C. At each position, a progressive (front-to-back) thrust optic flow stimulus is presented: first as square-wave grating, then as a starfield. In each pitch position, the same direction of thrust motion is presented, while the arena only shows the equatorial, north, or south portions of the full-field stimulus.

**Movie S3: Tethered flying fly with yaw and pitch control in a UV/green virtual environment, related to Figure 6, Figure S4**.

A tethered flying fly positioned inside a 9 column, 4 row G4 arena (270° partial cylinder) displaying a closed-loop virtual environment featuring a single dark vertical bar, checkerboard UV sky, and checkerboard green ground. The fly controls the visual scene’s rotational velocity through the detected wingbeat differential (L-R) and the pitch position relative to the horizon through the wingbeat amplitude sum (L+R), while the ground is constantly translating front-to-back to simulate the fly’s constant forward movement.

**Movie S4: Calculation of a 2D orientation histogram in a virtual environment with yaw and pitch control, related to Figure 6**.

(top) An “unwrapped” view of the cylindrical G4 arena while displaying a closed-loop virtual environment featuring a UV sky, green ground, dark green horizon, and single dark vertical stripe. As the fly modulates its wingbeat amplitude differential (L-R) and sum (L+R), the yaw rotational position of the stripe and the pitch position of the horizon are changed accordingly. (bottom) The 2D histogram of the fly’s yaw (x-axis) and pitch (y-axis) position, beginning from the trial start over the course of the trial. The center of the histogram represents the [0,0] orientation of the visual scene (when the vertical stripe is straight ahead, and the horizon is level). More time spent at any specific orientation results in a brighter color at its associated position in the histogram.

